# piqtree: A Python Package for Seamless Phylogenetic Inference with IQ-TREE

**DOI:** 10.1101/2025.07.13.664626

**Authors:** Robert Neil McArthur, Thomas King-Fung Wong, Yapeng Lang, Richard Andrew Morris, Katherine Caley, Vijini Mallawaarachchi, Bui Quang Minh, Gavin Huttley

## Abstract

`piqtree` (pronounced pie-cue-tree) is an easy to use, open-source Python package that provides Python script based control of IQ-TREE’s phylogenetic inference engine. `piqtree` builds IQ-TREE as a Python package, presenting a library of Python functions for performing many of IQ-TREE’s capabilities including phylogenetic reconstruction, ultrafast bootstrapping, branch length optimization, model selection, rapid neighbor-joining, alignment simulation, and more. As `piqtree` explicitly uses IQ-TREE’s phylogenetic algorithms, the computational and statistical performance of `piqtree` equal that of IQ-TREE. Modestly higher memory usage may be expected owing to the Python runtime and the need to load the alignment in Python. By exposing IQ-TREE’s algorithms within Python, `piqtree` offers users a greatly simplified experience in development of phylogenetic workflows through seamless interoperability with other Python libraries and tools mediated by the cogent3 package. It enables users to perform interactive phylogenetic analyses and visualization using, for instance, Jupyter notebooks. We present the key features available in the `piqtree` library and a small case study that showcases its interoperability. `piqtree` is distributed for use as a standard Python package at https://pypi.org/project/piqtree/, documentation is available at https://piqtree.readthedocs.io and source code at https://github.com/iqtree/piqtree.

## Introduction

We introduce `piqtree`, an open-source package available on all major operating systems, that offers a Python interface to the maximum-likelihood phylogenetic inference capabilities of IQ-TREE. Since the phylogenetic calculations are done by directly calling internal IQ-TREE routines, `piqtree` performance matches that of IQ-TREE (Table S1). By being implemented in Python, a widely used scripting language, `piqtree` supports interactive analyses and simplifies the integration of IQ-TREE into phylogenetic analyses via `cogent3` functions. Bespoke analyses that integrate IQ-TREE’s advanced algorithms can now be done with ease using Python scripts.

IQ-TREE (Nguyen et al., 2015; Minh et al., 2020; Wong et al., 2025) offers a command-line interface to phylogenetic inference using maximum likelihood. Beyond direct usage by researchers, IQ-TREE has been integrated into other applications and pipelines. These include the workflow platform Galaxy (The Galaxy Community, 2024), the phylogenetics gateway CIPRES (Miller et al., 2010), the microbiome analysis application QIIME2 (Bolyen et al., 2019), the orthology inference OrthoFinder (Emms and Kelly, 2019), and the real-time tracking of pathogen evolution Nextstrain (Hadfield et al., 2018).

The increase in large scale multi-gene phylogenomic data sets is requiring individual researchers to develop their own customized workflows that integrate phylogenetic reconstruction tools. At present, integrating phylogenetic packages involves using them as external applications. This requires implementing custom algorithms for controlling external applications, followed by processing of exported text file outputs (e.g. Hadfield et al., 2018). When the exported data file formats change, the analyses that depend on them can break, requiring work by the user to fix. In addition, this approach requires learning multiple programming languages (e.g. bash and Python), multiple software package managers (e.g. `homebrew, conda, pip`) and increases the challenge of ensuring the work is reproducible.

We have developed `piqtree` to directly expose the functionality of IQ-TREE in Python. Python is a highly popular programming language with an extensive ecosystem of scientific packages. `piqtree` is not intended as a replacement for the IQ-TREE application, but rather as a modular Python interface to its efficient algorithms. While not all of IQ-TREE’s functionality is currently available in `piqtree`, the overlap is substantial and growing with input from the user community.

The integration of IQ-TREE into Python is mediated through `cogent3` (COmparative GENomics Toolkit 3, Huttley et al., 2025a), the successor to PyCogent (Knight et al., 2007). `cogent3` provides extensive capabilities for data handling, data visualization, and robust algorithms for molecular evolution (Schranz et al., 2008; Lin et al., 2012; Wheeler et al., 2025). It implements unique substitution models, including context-dependent, non-reversible, and non-stationary models (e.g. Yap et al., 2010; Verbyla et al., 2013; Kaehler et al., 2015, 2017; McShea et al., 2025), and is the only software available for computing branch lengths when the substitution process is non-stationary (Kaehler et al., 2015). In `piqtree`, however, its primary role is to provide a library of fundamental biological data objects and to simplify integration of `piqtree` with other tools.

We introduce the key features of `piqtree` and present a small case study emphasizing the ease of developing a novel phylogenetic analysis that exploits data-level parallelism (described below) using the `cogent3` app ecosystem. `piqtree` can be installed from the Python Package Index. The data and code used for this work is available at DOI 10.5281/zenodo.15875241. An archive of the `piqtree` version is DOI 10.5281/zenodo.15860648. The source code and documentation are available at https://github.com/iqtree/piqtree and https://piqtree.readthedocs.io respectively.

## Key Features

### Computational performance

As `piqtree` uses IQ-TREE for all phylogenetic calculations, the time taken to execute an analysis is indistinguishable. Because communicating from Python to the IQ-TREE library requires in-memory copying of alignment data, `piqtree` memory usage is a modest increase over that of IQ-TREE (see Table S1).

### Phylogenetic reconstruction

A maximum-likelihood phylogenetic tree can be constructed from a `cogent3` alignment object under a chosen substitution model with the `build_tree` function. Users can further customize the model by specifying how state frequencies are chosen (either empirical, equal, or optimized through maximum likelihood). Further, rate heterogeneity across sites can be incorporated by allowing for a proportion of invariable sites and by using the discrete Gamma or FreeRate models with a custom number of rate categories (Yang, 1994b; Gu et al., 1995; Yang, 1995; Soubrier et al., 2012). Ultrafast bootstrapping can be performed by specifying the number of bootstrap replicates (Minh et al., 2013; Hoang et al., 2018). The number of threads to parallelize across can be specified for more efficient computation, and a random seed may be set for reproducibility. `piqtree` currently supports all IQ-TREE DNA models (including Lie Markov models, Woodhams et al., 2015), and amino-acid exchange rate matrices. The full listing of the available models can be found in the documentation.

### Fitting branch lengths to a tree topology

The `fit_tree` function enables branch lengths to be fitted to a fixed tree topology for a `cogent3` alignment object and a chosen substitution model. The function supports the same models as `build_tree`. The number of threads to parallelize across may also be set for faster computation. Nucleotide or amino acid probabilities and rate parameters are annotated as appropriate along the resulting `cogent3` tree object’s edges.

### Simulating an alignment

An alignment can be simulated over a `cogent3` tree object for a chosen model with AliSim (Ly-Trong et al., 2022, 2023) using the `simulate_alignment` function. The sequence length, root sequence for the simulated alignment, as well as insertion and deletion rates and their respective size distributions can be specified. A random seed may be chosen for reproducibility and the number of threads for parallelization can be specified for faster computation.

### Selecting an evolutionary model

The `model_finder` function exposes the model finding capabilities of IQ-TREE. The resulting object provides convenient interfaces to the models selected by the different information measures. The model selected according to the user preferred information criteria (e.g. BIC or AIC, Posada and Buckley, 2004) can be easily applied to either `build_tree` or `fit_tree`.

### Estimating pairwise distances

The `jc_distances` function computes pairwise genetic distances from a `cogent3` alignment object using the JC69 model (Jukes and Cantor, 1969), returning a `cogent3` distance matrix object. The user can specify the number of threads to use for the computations.

### Estimate a phylogeny using RapidNJ

From a `cogent3` distance matrix object, the nj_tree function uses the efficient Rapid Neighbour-Joining tree algorithm used internally by IQ-TREE (Simonsen et al., 2011). When a user has `piqtree` installed, methods on `cogent3` alignments and distance matrix objects will use this implementation rather than the native `cogent3` methods. By default, negative branch lengths resulting from the method are set to 0.

### Simulate a random tree

The `random_tree` function can be used to make a randomly generated tree. The number of taxa and strategy used to generate the tree can be specified. Options to generate include balanced trees, birth-death trees, caterpillar trees, star trees, uniform trees, or Yule-Harding trees. A random seed can be specified to ensure reproducibility.

### Constructing a consensus tree

The `consensus_tree` function can be used to construct a consensus tree from a collection of trees on the same taxa set. By default, the majority-rule consensus tree is constructed (Margush and McMorris, 1981). Additionally, a minimum support value (0-1) can be specified, representing a proportion of trees. Consequently, a clade must appear in more trees than this proportion in order to appear in the resulting consensus tree. If a minimum support of 1 is specified, the algorithm constructs the strict consensus tree (Sokal and Rohlf, 1981). If a minimum support of 0 is specified, the algorithm constructs an extended majority-rule consensus tree (Felsenstein, 1985).

## Case study – demonstrating a novel phylogenetic analysis by combining piqtree with other Python packages

In this case study we use a supertree method to estimate the root of a phylogeny from phylogenomic data, an analysis not natively supported by IQ-TREE. We demonstrate how a user can perform this analysis by combining existing Python tools with `piqtree`.

Supertree methods produce a consensus tree from a collection of trees that have overlapping, but not identical, taxa sets. The Python package `sc_supertree` (McArthur et al., 2024) requires rooted trees as input. `piqtree` implements non-reversible substitution models that can generate rooted phylogenies using statistical estimation of the root position (see Yap and Speed, 2005), i.e., no prior knowledge about an outgroup lineage is required.

### The data

We selected the published phylogenomic data set of Chiari et al. (2012) for the case study. The data consisted of 248 one-to-one orthologous protein-coding genes from turtles, crocodilians, and birds, with the lungfish included as an outgroup. This data set was selected as it satisfied the requirements of the supertree method: not all genes were present in all taxa, and nucleotide data, required by the UNREST model (Yang, 1994a), was available. The original data set was a single concatenated alignment derived from alignments of the 248 genes. A partition file, delineating the gene boundaries in the concatenated alignment was available.

The goal of the original study was to assess the relative positions of turtles, crocodilians, and birds using the Lungfish as a outgroup. Chiari et al. (2012) used RAxML (Stamatakis, 2006) with time-reversible nucleotide and amino substitution models along with a range of rate heterogeneity strategies (Chiari et al., 2012).

We emphasize that our intent with selecting this data was not to present the following analysis as necessarily superior to that employed by Chiari et al. (2012).

### Preparing the data

We used data handling features of `cogent3` to split the concatenated alignment into alignments of the original 248 genes. In the concatenated alignment, where an ortholog was missing from a species, it was represented by a sequence consisting entirely of ‘?’. We omitted species from the alignment in these cases. The outcome of this data preparation (see Figure S2 for this algorithm) was 248 fasta formatted alignment files that were the input for our analysis.

### Combining cogent3, piqtree and sc_supertree to statistically estimate a root to the tetrapod tree

A common feature of phylogenomic studies is the need to analyze many separate sequence alignments, a process well suited to data-level parallelization. By distributing different analyses across multiple CPUs and running them simultaneously, computation times can be greatly reduced. For instance, if two alignments are processed one after the other on a single CPU, the total time equals the sum of their individual processing times. However, if each alignment is processed simultaneously on a different CPU, the time can be cut by as much as half, illustrating the benefits of parallel processing.

To produce a phylogeny from an alignment file, we load the file into memory using `cogent3` and pass the resulting alignment to `piqtree`, which returns a phylogeny. Since we are analyzing individual alignments, and some may be too short to provide reliable estimates, we eliminate alignments shorter than 471 bases. This threshold was chosen because it excludes the shortest 5% of alignments and is divisible by 3 so each codon position is equally represented in the result. With a directory containing 248 FASTA-formatted multiple sequence alignment files, we perform this procedure for each file, collecting the resulting phylogenies. We then pass this collection to `sc_supertree`, which generates a single consensus supertree. To expedite phylogeny construction, we use data-level parallelization. In this setup, `piqtree` is responsible only for estimating a phylogeny from a single file. The selection, loading of FASTA files and filtering by length is managed by `cogent3`, and the final consensus step is performed by `sc_supertree` (McArthur et al., 2024).

`cogent3` provides an “app” mechanism to validate the interoperability of different functions. An app is a single-purpose Python function that specifies input and return data types. `cogent3` uses these definitions to enable multiple apps to be combined into a “composed pipeline” that efficiently utilizes data-level parallelism. `cogent3`’s data loading functions track the source files for each object, allowing apps to automatically track the provenance of every result during execution. The built-in `cogent3` apps are based on standard `cogent3` functions, which users can also use in traditional algorithms.

`piqtree` has exposed many IQ-TREE functions as apps to take advantage of `cogent3` app capabilities. All app names take the form `piq_<function name>`, for example the app for `build_tree` is named `piq_build_tree`. All `piqtree` functions and apps record the provenance of generated output. e.g. tree objects returned by `piqtree` have attributes that identify the path to the alignment file that produced it along with model settings.

To solve the analysis here, we present algorithms for using apps or using standard functions. In Figure 1, we use `cogent3` apps to simplify the expression of finding fasta formatted files, filtering alignments by length, looping and distributing computations across CPUs. In Figure 2, we express the problem using conventional functions without parallel execution. In both cases, we specify random number seeds for the `piqtree` and `sc_supertree` algorithms to ensure reproducibility.

**Fig. 1:**
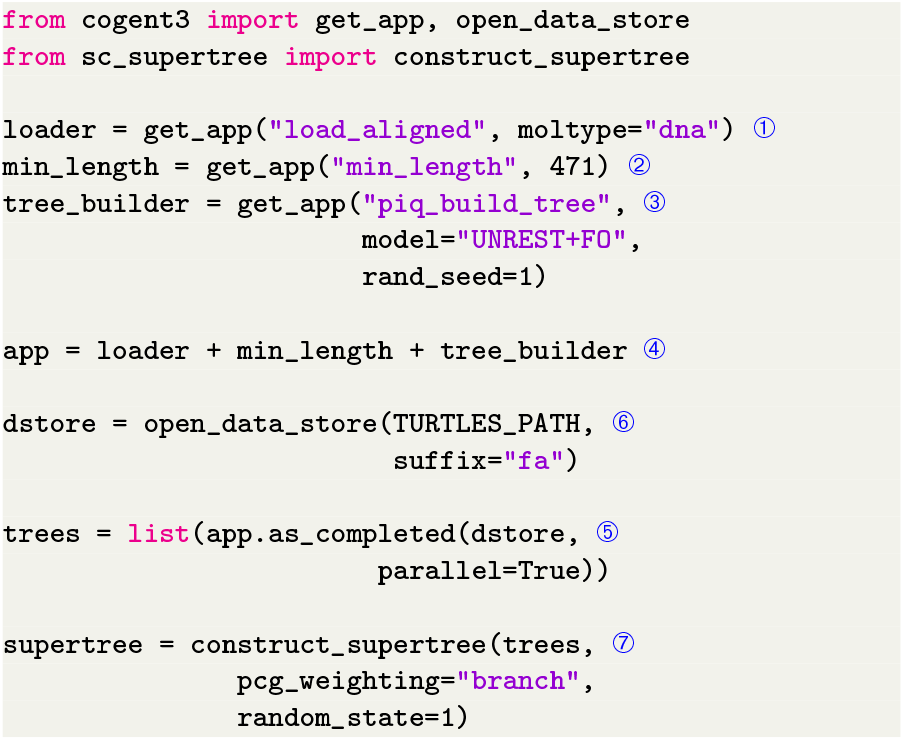
Combining `piqtree` with other apps to construct a rooted supertree. A sequence loader ➀, a filter for alignments with length *≥* 471 ➁ and `piqtree` tree builder using the UNREST+FO model ➂ are added together to produce a composed app ➃ and applied in parallel ➄ to a directory (`TURTLES_PATH`) of alignment files (file names end with .fa) ➅ to produce a list of `cogent3` tree objects. Spectral Cluster Supertree ➆ is then applied to construct the supertree from these trees. The rand_seed and `random_state` arguments are set to a specific value to ensure reproducibility. (See the script `piqtree_app_demo.py` in DOI 10.5281/zenodo.15875241 for the functioning version.)

**Fig. 2:**
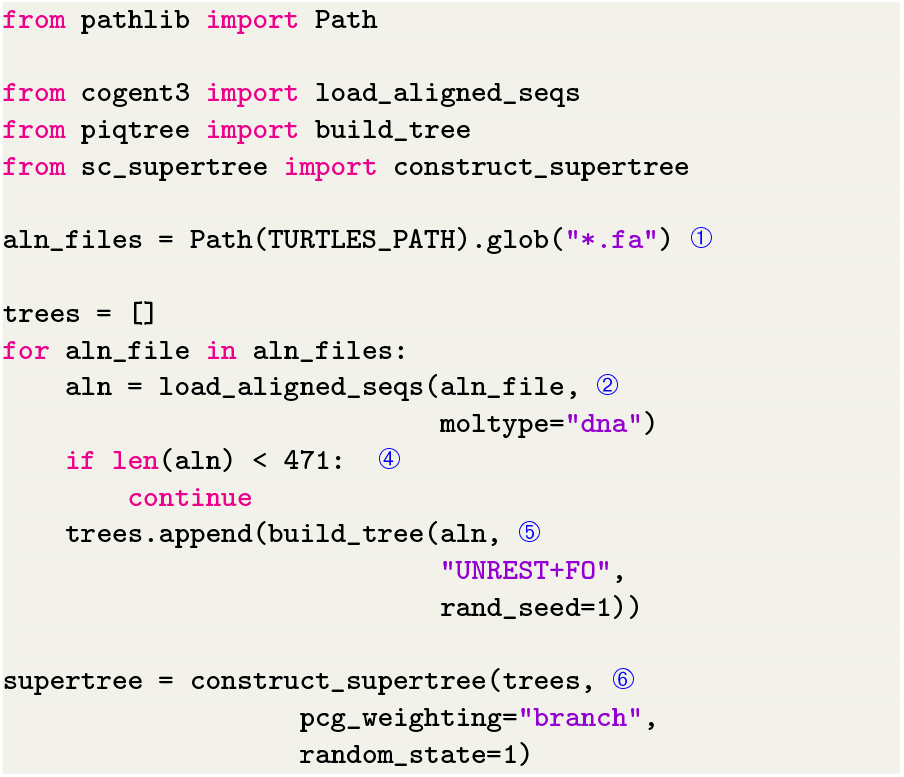
Combining `piqtree` with other tools using standard functions to construct a rooted supertree. The FASTA files for each gene are extracted from the data directory ➀. Each alignment is loaded into Python ➁ and checked to ensure the alignment length is *≥* 471 before `build_tree` is used to reconstruct the corresponding gene tree under the UNREST+FO model ➂. Spectral Cluster Supertree ➃ is then applied to construct the supertree from these gene trees. There is no parallelization of the computations. (See the `piqtree`_demo.py script in DOI 10.5281/zenodo.15875241 for the functioning version.)

The algorithms presented in the figures are selected for brevity rather than as general solutions for all phylogenomic studies. For example, consider a case where the number of species is much larger and the time taken to estimate the phylogeny substantially longer. In such instances, it is better practice to write the generated phylogenies to disk to avoid recalculation should the program need to be restarted. `cogent3` provides writer apps for this purpose, whose usage is described in the `cogent3` documentation. These apps also use automated logging of analyses, tracking the provenance of all results and thus facilitating computational reproducibility. (See the `piqtree_app_data_store_demo.py` script in DOI 10.5281/zenodo.15875241 for a demonstration for the case study.)

The result of the analysis (Figure 3) exactly recapitulates the Chiari et al. (2012) tree from nucleotide analysis with the exception of the root placement. The statistical estimate of the root in the supertree is two edges away from the expected position on the Protopterus (lungfish) branch. As it is long established that stationary non-reversible models have low statistical power to detect the root position (Yap and Speed, 2005) this result is not unexpected.

**Fig. 3:**
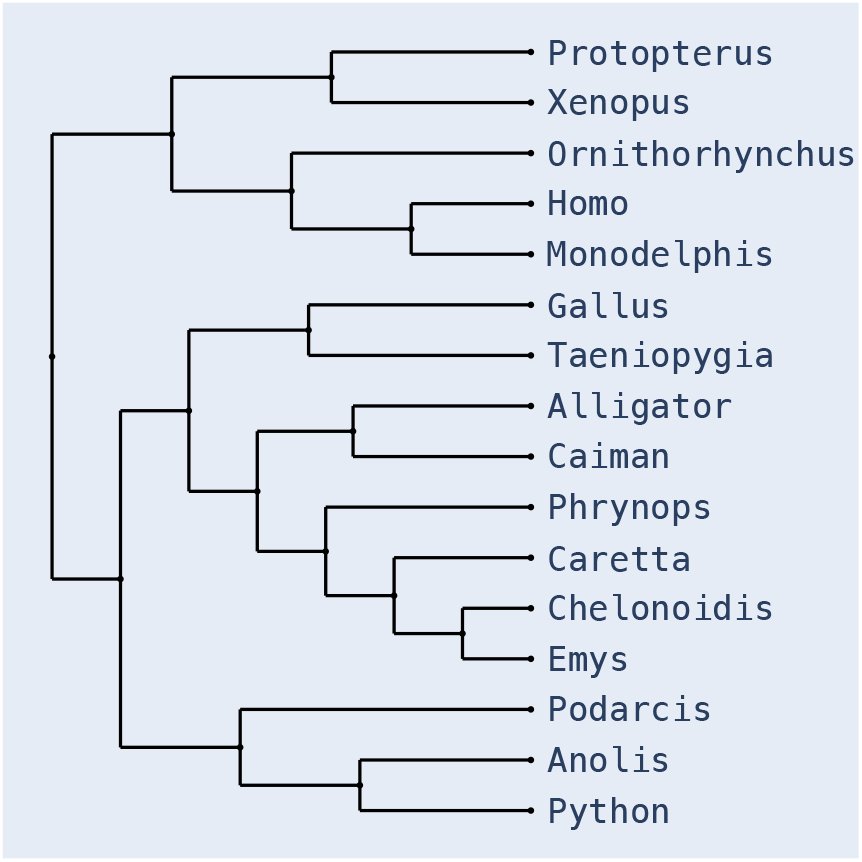
The supertree formed from the gene trees of the turtles dataset. The image was created using the `cogent3` built-in dendrogram algorithms which use the Plotly interactive graphics library (Plotly Technologies Inc, 2015).

We caution readers against placing too much emphasis on the exactitude of the presented results. For this analysis, we found the produced supertree was sensitive to the choice of random number seeds. There were two topologies produced: the one shown in Figure 3, or an alternate in which the two lizards (Anolis and Podarcis) were paired. In the original tree, this node had the weakest bootstrap support (53%, Chiari et al., 2012, Fig 3b).

### The phylogenetic Python data science ecosystem

Users of `piqtree` will be able to take advantage of Python’s sophisticated support for interactive exploratory data analyses and other Python phylogenetic packages that complement it. For instance, while `cogent3` has its own built-in tree visualization capabilities (e.g. Figure 3) the ETE3 toolkit provides more comprehensive capabilities (Huerta-Cepas et al., 2016) for interactive editing. A `cogent3` app cogent3-ete3 can be used to transform the `cogent3` trees into ETE3. iplotx (Zanini, 2025) provides a sophisticated alternative for producing static images and has conversion routines for `cogent3` tree objects. The `diverse-seq` package (Huttley et al., 2025b) provides `cogent3` apps to facilitate development of phylogenetic workflows. Phylogenetic algorithms, from sequence alignment to phylogenetic reconstruction itself, all scale poorly with the number of sequences. From a large collection of sequences, `diverse-seq` selects a user nominated number as representative of the sequence diversity in the collection. By reducing the number of sequences and preserving the diversity, it reduces the computational resources needed to build a workflow prototype. The `phylim` package (Lang and Huttley, 2025) provides `cogent3` apps and works with `piqtree` to check that a fitted phylogenetic model is valid. Phylim draws on mathematical results concerning the uniqueness of parameter estimates from a fitted phylogenetic models (Chang, 1996; Kaehler, 2017). It evaluates whether the fitted parameters satisfy the mathematical conditions required for a unique solution, addressing uncertainties regarding the reliability of inference when site-saturation is a concern.

## Conclusion

`piqtree` combines the algorithmic sophistication of IQ-TREE with the accessibility and flexibility of data analysis using Python. While the two tools share the same underlying algorithms, `piqtree` offers distinct advantages over IQ-TREE in certain contexts. For workflow developers, it provides a consistent installation process across operating systems and a stable application programming interface that will not break between major releases of IQ-TREE. For users seeking to undertake interactive analyses, `piqtree` provides a rich user experience in computational environments such as Jupyter Notebooks. By being implemented in Python, `piqtree` makes adaptation of a phylogenetic inference engine to novel applications possible by the users themselves without requiring input from the IQ-TREE development team. At the time of writing, there were 459 packages listed at the Python Package Index (https://pypi.org) that matched the search term “phylogenetics”. As we have shown with `sc_supertree`, `piqtree` and `cogent3` can simplify interoperability with other tools, reducing a barrier to innovation in phylogenetic analyses. As its adoption by other bioinformatic tools (Mallawaarachchi et al., 2023) highlights, `piqtree` can increase development of new phylogeny based methodologies.

## Acknowledgments

Chan-Zuckerberg Initiative EOSS5-0000000223 awarded to GAH, VM and BQM and EOSS4-0000000312 to BQM. National Science Foundation 2333243 awarded to Joanna Masel with a subcontract to BQM.

