## Supplementary Material for "piqtree: A Python Package for Seamless Phylogenetic Inference with IQ-TREE"

### Performance comparison between `piqtree` and IQ-TREE

We compared the computational performance of `piqtree` with IQ-TREE. Both were compiled in the same runtime environment. We compared the time needed to reconstruct a phylogenetic tree under a GTR+G model for an alignment of length 10000 of 500 taxa. The alignment was selected purely for its large size making it suitable for benchmarking, and was the first simulated alignment with 500 taxa from the SCS dataset (McArthur et al., 2024). The same seed was used over five trials, and a single thread was used. The minimum, mean and maximum times are reported in Table S1. Both `piqtree` and IQ-TREE solve the problem in the same amount of time. There is no discernible overhead that comes from marshalling the data from Python to IQ-TREE's internal representation. Higher memory usage can be expected with `piqtree` as the alignment is stored in memory in both Python and IQ-TREE, plus overhead from the Python runtime environment.

| | Min | Mean | Max |
|---|---|---|---|
| `piqtree` Time | 1350.28s | 1357.87s | 1369.33s |
| IQ-TREE Time | 1363.69s | 1375.07s | 1387.82s |
| `piqtree` Memory | 1111.37 MiB | 1112.53 MiB | 1113.04 MiB |
| IQ-TREE Memory | 864.64 MiB | 865.31 MiB | 867.32 MiB |

Table S1: `piqtree` and IQ-TREE compute times were approximately equal. `piqtree` uses more memory to load and store the alignment and Python runtime.

### Selected Algorithms

```python
def build_iqtree(aln_file, out_file, substitution_model="GTR", nthreads=1,
↪ tree_builder_args=None):
    """
    build tree using piqtree
    arguments:
        aln_file    file name of input aligment
        out_file    file name to write tree to
    """
    import cogent3 as c3

    aln = c3.load_aligned_seqs(aln_file, moltype="dna")
    seqname_aliases = {n: f"s{i}" for i, n in enumerate(aln.names)}
    rn_aln = aln.rename_seqs(lambda x: seqname_aliases[x])
    build_tree = c3.get_app(
        "piq_build_tree",
        model=substitution_model,
        num_threads=nthreads,
        optional_args=tree_builder_args,
    )
    tree = build_tree(rn_aln)
    rn_tree = tree.renamed_nodes({a: n for n, a in seqname_aliases.items()})
    rn_tree.write(out_file)
    return rn_tree, aln.names
```

Figure S1: Updating Nextstrain to use `piqtree` and `cogent3`. The original Nextstrain algorithm was 109 lines of code.

```python
import pathlib
from cogent3 import load_aligned_seqs

def parse_nexus_charsets(partition_path: pathlib.Path) -> dict[str, tuple[int, int]]:
    """return the partition name, alignment start and stop from a nexus file."""
    result = {}
    for line in partition_path.read_text().splitlines():
        line = line.strip()
        if not line.lower().startswith("charset"):
            continue

        _, name, _, start, _, stop = line[:-1].split()
        # convert 1-based coordinates to 0-based
        result[name.split(".")[0]] = (int(start.strip()) - 1, int(stop.strip()))
    return result

# turtle.nex and partition.nex originally from Chiari et al
aln = load_aligned_seqs("turtle.nex", moltype="dna")
# partition nex defines the boundaries for each gene
partitions = parse_nexus_charsets(pathlib.Path("partition.nex"))

outdir = pathlib.Path("turtle_partitions")
outdir.mkdir(exist_ok=True)
for gene_name, (start, stop) in partitions.items():
    gene_aln = aln[start:stop]
    # a 'length' is the number of non-gap characters in a sequence
    lengths = gene_aln.get_lengths()
    # only keep sequences in an alignment if they have nucleotide data
    gene_aln = gene_aln.take_seqs_if(
        lambda seq: lengths[seq.name] > 0,
    )
    outpath = outdir / f"{gene_name}.fa"
    gene_aln.write(outpath)
```

Figure S2: Extracting genes from concatenated Chiari et al alignment

```python
import cogent3 as c3

aln = c3.get_dataset("brca1")
dmat = aln.distance_matrix(calc="TN93")
tree = dmat.quick_tree()
```

Figure S3: Combining features of `cogent3` and `piqtree`. The TN93 genetic distance is not provided by `piqtree`, so `cogent3` will be used for distance calculation. If `piqtree` is installed, then its rapidNJ algorithm will be used by `quick_tree()` for phylogeny estimation.
